# A molecular model of the surface-assisted protein aggregation process

**DOI:** 10.1101/415703

**Authors:** Y.G. Pan, S. Banerjee, K. Zagorski, L.S. Shlyakhtenko, A.B. Kolomeisky, Y. L. Lyubchenko

## Abstract

The importance of cell surfaces in the self-assembly of proteins is widely accepted. One biologically significant event is the assembly of amyloidogenic proteins into aggregates, which leads to neurodegenerative disorders like Alzheimer’s and Parkinson’s. The interaction of amyloidogenic proteins with cellular membranes appears to dramatically facilitate the aggregation process. Recent findings indicate that, in the presence of surfaces, aggregation occurs at physiologically low concentrations, suggesting interaction with surfaces plays a critical role in the disease-prone aggregation process. However, the molecular mechanisms behind on-surface aggregation remain unclear. Here we provide a theoretical model that offers a molecular explanation. According to this model, monomers transiently immobilized to surfaces increase the local monomer protein concentration and thus work as nuclei to dramatically accelerate the entire aggregation process. This theory was verified by experimental studies, using mica surfaces, to examine the aggregation kinetics of amyloidogenic-synuclein protein (α-Syn) and non-amyloidogenic cytosine deaminase APOBEC3G (A3G).

## Main text

The assembly of proteins into aggregates of various type is a general phenomenon found frequently in both natural and industrial processes (1, 2). Different types of protein aggregates are commonly observed. For example, proteins can self-assemble into filamentous aggregates; the actin filament is one of the numerous examples of this process. Another, and the most known example, is the formation of aggregates by amyloidogenic proteins. According to the current views, the formation of amyloidogenic aggregates is a hallmark in the development of numerous disorders, including neurodegenerative diseases such as Alzheimer’s disease (3). Although the self-assembly of protein aggregates can take place in a solution, the importance of membrane surfaces in such processes is also acknowledged (e.g., (4-6)).

In general, the aggregation process is accelerated in the presence of membranes (e.g., (7) and references therein). With respect to Alzheimer’s disease, great interest has been given to the role of membranes in disease pathogenesis and in facilitating the assembly of amyloid fibrils (*e.g.,* reviews (6, 8-12)). Importantly, the inclusion of cholesterol and gangliosides into membranes changes the structure and stability of amyloid agregates; these changes appear to contribute to the neurotoxic effect of aggregates (10), (11). However, a molecular mechanism explaining the catalysis of membrane surfaces toward protein aggregation remains poorly understood.

Recent studies have shown that, similar to membranes, surfaces such as glass (13), mica (14, 15), and zeolites (16) also accelerate the aggregation process for various amyloidogenic proteins. Importantly, amyloid aggregates, primarily fibrils, have been imaged using electron microscopy (16) and AFM (14, 15). As such, the use of solid surfaces has made it possible to partially visualize the molecular mechanism behind the surface-acceleration effect. According to the model proposed in (14, 15), the accelerated aggregation is due to the fast, two-dimensional diffusion of amyloid peptide molecules at the surface-liquid interface.

AFM has been used to directly observe the accelerated aggregation of amyloid peptides and the alpha-synuclein (α-Syn) protein on mica surfaces. Results demonstrated that the assembly of proteins into aggregates took place at a low protein concentrations, while no aggregation was detected in bulk solution (17). Importantly, time-lapse AFM experiments in liquid did not reveal fast mobility of molecules at the mica-liquid interface (17). These data were also in line with observations performed using time-lapse high-speed AFM (18, 19). These results suggest an alternative mechanism of accelerated aggregation on the surface that does not include surface diffusion. Such a mechanism is proposed in the current work.

Here a novel theoretical model is provided to explain the molecular mechanism of the surface-mediated catalysis behind the protein aggregation process. According to this model, aggregation starts with protein monomers transiently attaching to the surface due to molecular interactions. This process increases the local concentration of proteins, which in turn increases the probability of oligomerization reactions to occur on the surface. Based on this model, aggregation occurs by the assembly of oligomers on these transiently bound monomers. This theoretical prediction was experimentally tested using two proteins that follow different aggregation pathways. One such protein, α-Syn, is a typical amyloidogenic protein capable of assembling into aggregates of various morphologies, including fibrils. The other protein, cytosine deaminase APOBEC3G (A3G), assembles into oligomers of various sizes depending on the protein concentration (e.g., (20) and references therein).

To explain the complex processes of surface-assisted protein aggregation, a new theoretical model was further developed. The main assumptions for this theoretical model are as follows:

1. 1) Due to intrinsic interactions with surfaces, protein monomers very quickly bind to the surface and establish an effective equilibrium between surface-bound and free monomers in solution. The equilibrium coverage is given by the parameter 0 < θ < 1, which describes what fraction of the surface is covered by protein monomers.
2. 2) The effective concentration of monomers near the surface increases in comparison to concentration in bulk solution, and this accelerates the rates of oligomerization.

To examine these arguments quantitatively, *C(t)* can be defined as the time-dependent concentration of protein monomers in solution. At *t = 0, C(t) = C*_*0*_. This represents the initial concentration of proteins in solution. Assuming that the surface area is equal to *L*^2^, and the molecular volume of one protein monomer is v_0_∼d^3^, the maximum possible number of monomers on the surface can be estimated as follows:

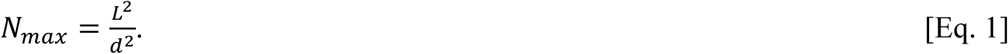

The number of adsorbed proteins can be given by *N*_*1*_, where

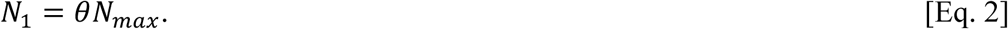

For a solution in the absence of a surface, the reaction rate for the formation of dimers as a first step of aggregation can be calculated as follows, where *k* is the bulk rate constant:

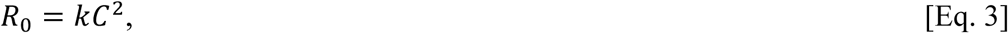

However, in the presence of the surface, the following equation can applied:

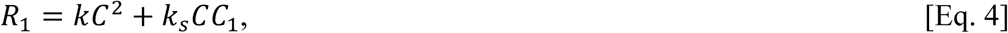

In the equation above, *k*_*s*_ represents the reaction rate for the formation of dimers on the surface, and *C*_*1*_ represents the concentration of the protein monomers in the volume around the surface.

It can be estimated using equations 1 and 2, with *N*_*A*_ being Avogadro’s number, as follows:

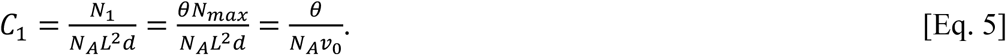

Now the acceleration factor can be evaluated in the reaction rate due to the presence of the surface at earlier times.

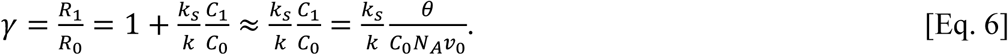

To estimate this factor using more or less realistic parameters, very low coverage is assumed, with θ = 0.001 (0.1%), C_0_ = 100 nM, *v*_*0*_ ∼ 100 nm^3^, and 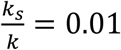. This is because the reaction rate on the surface is expected to be smaller than in bulk solution due to steric constraints and possible conformational changes. Next, γ ∼ 1.6*10^3^ can be obtained, which represents a markedly accelerated aggregation process due to the presence of the surface.

These calculations show that the aggregation in bulk solution is relatively slow; the dominating chemical process in the system is the formation of oligomers on the surface. This can be described by the following quasi-chemical reaction, where *C* represents bulk protein monomers; *C*_1_ represents surface-bound protein monomers; *C*_2_ represents surface-bound dimers:

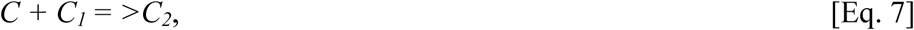

Next, *C*_*2*_*(t)* can be given as the time-dependent concentration of dimers, with *C*_*2*_*(t=0) =0* and the rate constant *k*_*s*_. As such, the chemical kinetic equations for this system can be written as follows:

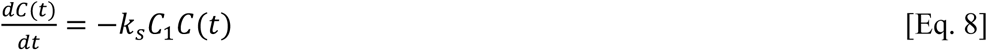

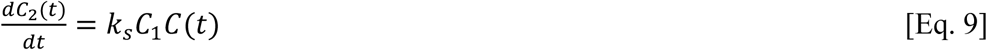

These equations can be easily solved to produce the following:

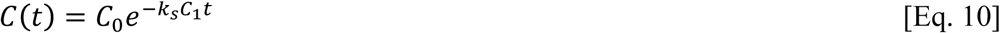

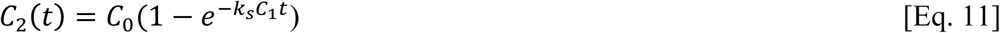

If the reaction rate on the surface is assumed to be relatively slow and/or the time relatively short, then equation 11 can be expanded to linear terms of time, yielding the following:

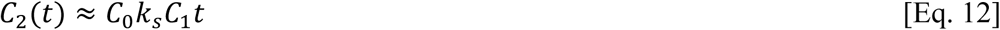

From this expression, the ratio of dimers-to-monomers on the surface as a function of time can be given by the following:

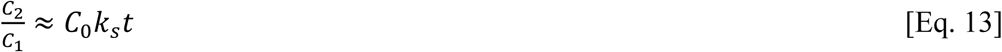

This result predicts that the ratio will grow linearly with time, and it will be proportional to the original concentration of the protein monomers in bulk solution.

To experimentally test these theoretical predictions, an approach developed by us recently (17) was applied, as shown schematically in Figure 1. A mica sheet was placed in a test tube (Fig. 1A) containing the protein solution and incubated at room temperature for finite time; afterwards, to directly count the number of aggregates appearing on the surface, the mica was removed, rinsed with water, dried, and imaged using AFM (Fig. 1B). Such experiments were performed at different incubation times. As a control, aliquots were taken from the same protein solution, but no mica strip was added (Fig. 1C). The AFM image for the control is shown in Fig. 1D.

**Figure 1.**
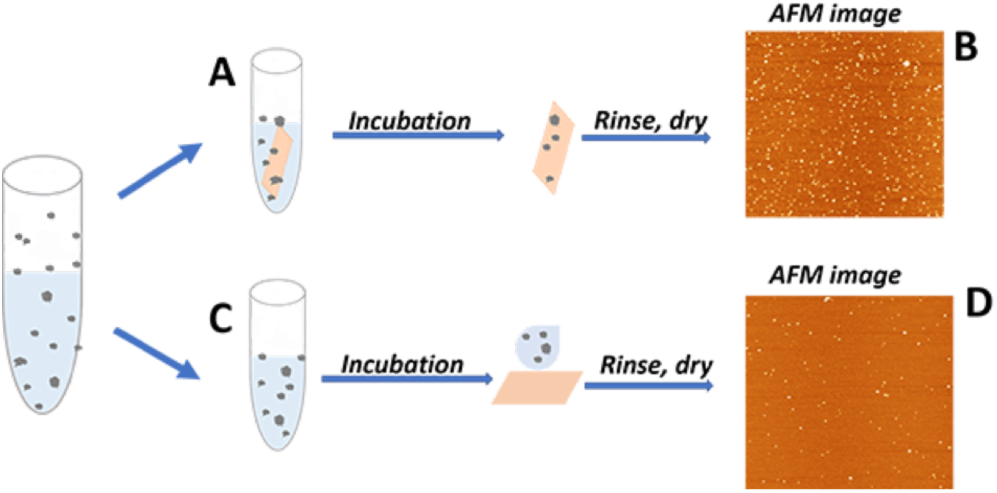
A schematic presentation of the experimental setup to monitor **(A)** the on-surface A3G aggregation, and in-parallel, **(B)** control experiments for aggregation in bulk solution. **C** and **D** are AFM images of on-surface and in bulk aggregation respectively.

Figure 2A shows the results of experiments with 2 nM α-Syn in the presence of mica, imaged between 0 and 26 hours. The data show that, over time, more globular features appear on surfaces, and their sizes also increase upon incubation. Supplemental Fig. S1 provides more images at times ranging from 1 hour to 20 hours. Control samples (Figs. 2B and S2) did not show an observable change in the number of aggregates. Figure S3 provides quantitative analysis of the AFM images; in this graph, the ratio of oligomers-to-monomers is plotted as a function of time. The value grows over time, reaching a plateau around 48 hours. The initial part of the kinetics of aggregation (between 0 and 20 hours) are fitted by a linear plot (Fig. 2C), supporting the prediction from the theoretical model in Eq. 13. The data for control experiments are shown in Fig. 2D and did not reveal aggregate assembly in bulk solution.

**Figure 2.**
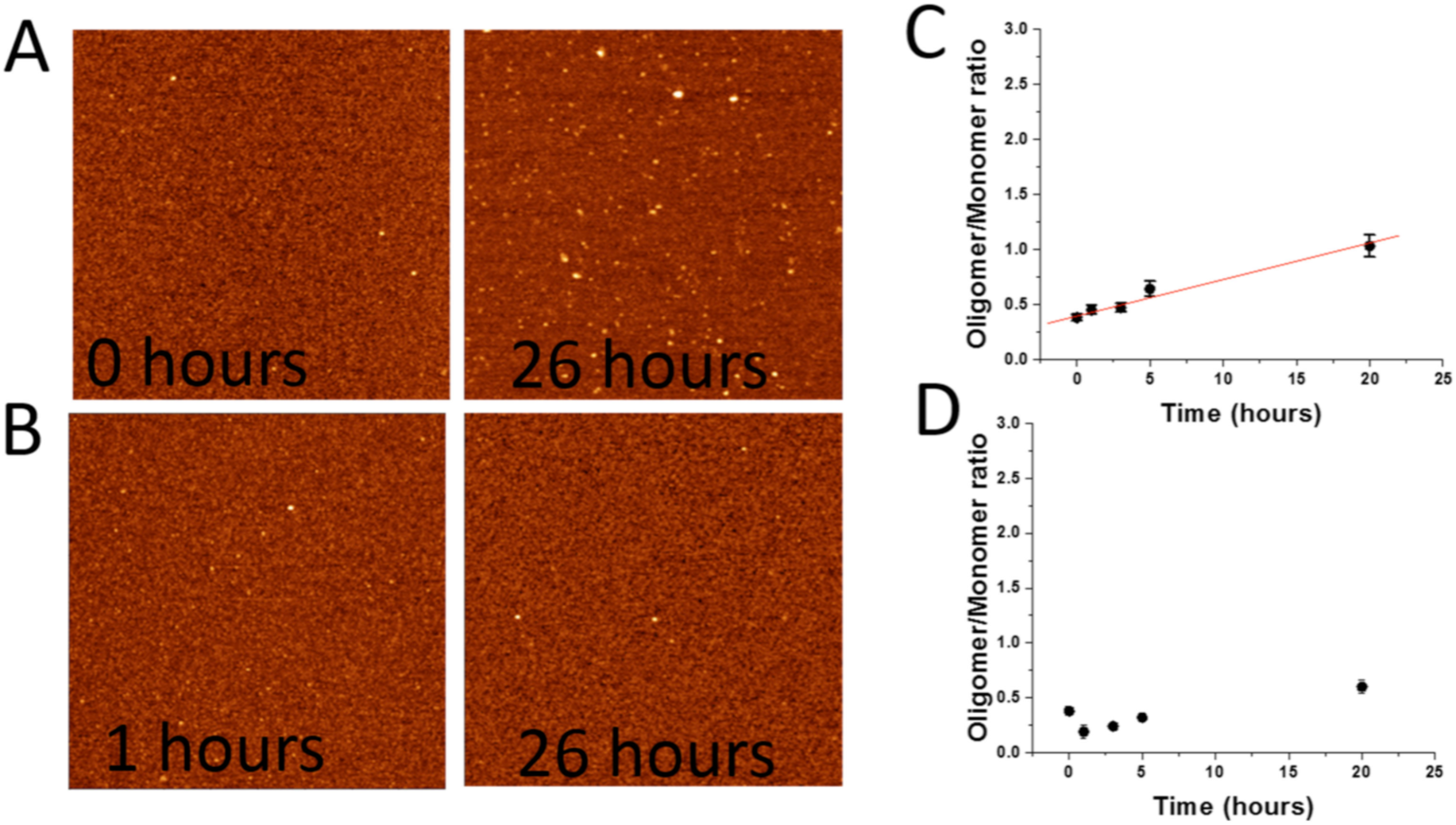
Experimental data for the aggregation of α-Syn. (**A**) AFM images of the on-surface aggregation of 2nM α-Syn for 0 and 26 hours. (**B**) AFM images of aggregation for 2nM α-Syn in the bulk solution for 1 hour and 26 hours. Scan sizes are 800 nm. (**C**) The time dependence of the oligomer-to-monomer ratio of 2nM α-Syn in the on-surface aggregation and (**D**) in bulk solution.

Similar experiments were performed using an increased concentration of α-Syn (10 nM). AFM images of the on-surface aggregation process, taken at different times during incubation, are shown in Fig. S4. These images clearly show that aggregates appear upon incubation, and their number and sizes increase over time. Control experiments obtained for the same incubation times (Fig. S5) do not show such time-dependent α-Syn aggregation in bulk solution. Quantitative analyses of these data for on-surface aggregation and in bulk solution are presented in Figs. S6A and S6B, respectively.

Similar to the data obtained for 2 nM α-Syn, the ratio of the number of oligomers-to-monomers increases gradually over time. The early aggregation kinetic graphs were fitted with the linear plot (Fig. S6A), and the slope for 10 nM of α-Syn turned out to be larger than for 2 nM α-Syn, which is in line with the theoretical predictions. Meanwhile, no time-dependent aggregation was observed for 10 nM α-Syn in the bulk solution (Fig. S6B), similar to control experiments for 2 nM of α-Syn.

To further test theoretical predictions, experiments were performed with the A3G protein, which has a strong propensity toward oligomerization depending on its concentration in solution (20). This feature is considered an additional mechanism for the antiviral activity of A3G. Figures 3A and 3B show AFM images of the on-surface aggregation of A3G at a concentration of 1 nM; the figures correspond to 5 min and 120 min and in the bulk solution, respectively. Figures S7A and S7B provide additional AFM images at intermediate time intervals for the aggregation of A3G in the presence of mica surface and in bulk solution, respectively. These images clearly show the accumulation of on-surface aggregates. No aggregation is observed for control experiments in bulk solution performed in parallel. Figure S8 provides a quantitative analysis of the aggregation data and demonstrates that the oligomer-to-monomer ratio increases over time. Figure 3C shows the initial process of the aggregation kinetics (between 0 and 2 hours) fitted with a linear plot. On the other hand, aggregation of A3G in solution is not time-dependent, showing no change in the oligomer-to-monomer ratio over time (Fig. 3D). Table S1 in the supporting information section assembles data characterizing the kinetics for the on-surface aggregation of both α-Syn and A3G proteins.

**Figure 3.**
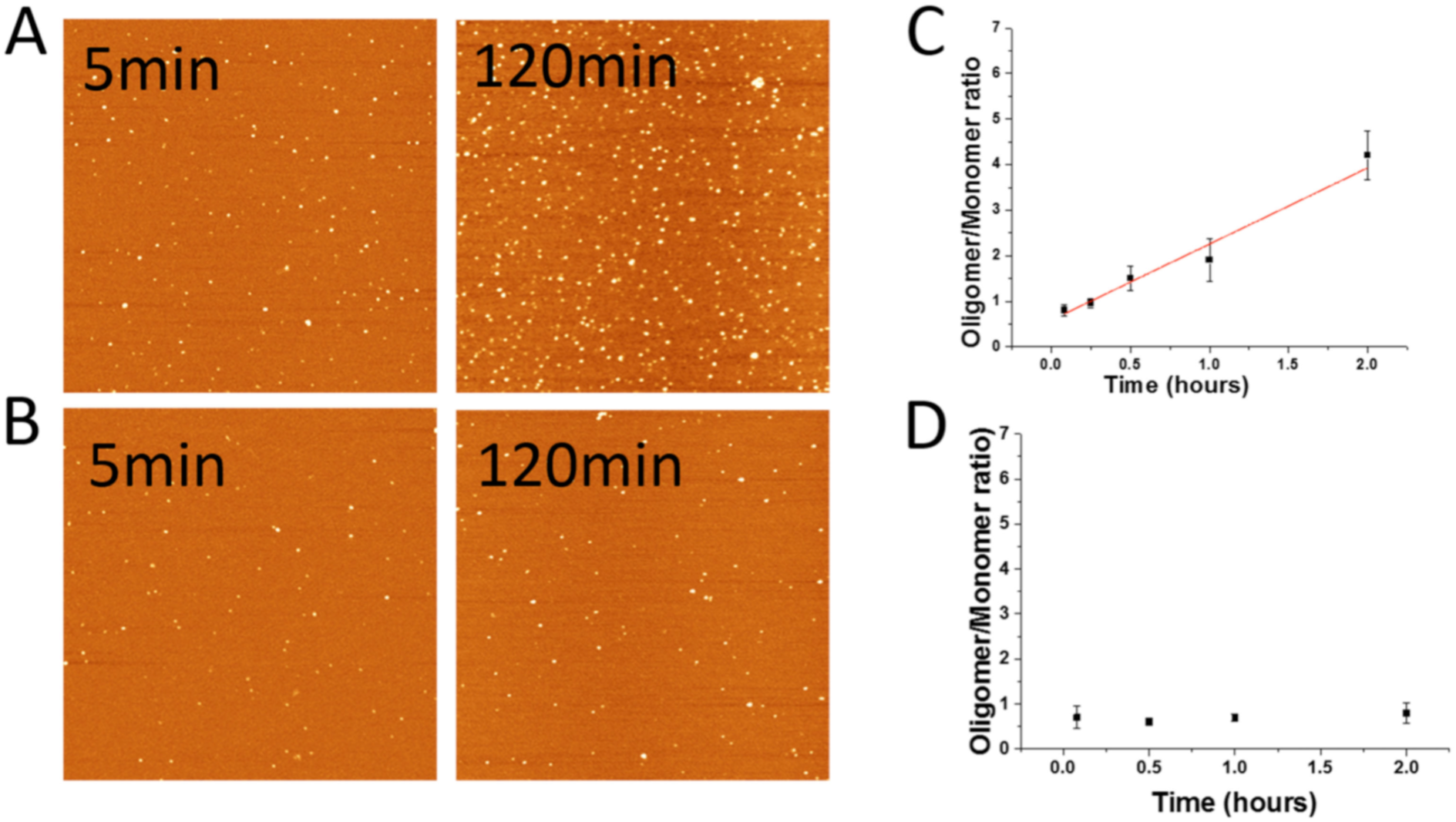
Experimental data for A3G protein aggregation. **(A)** AFM images of the on-surface aggregation of 1nM A3G at 5 min and 120 min. **(B)** AFM images of aggregation of 1 nM A3G in bulk solution at 5 min and 120 min. Scan sizes are 1.5 microns. **(C)** The time dependence of the oligomer-to-monomer ratio of 1nM A3G in the on-surface aggregation and **(D)** in the bulk solution.

Overall, the data obtained support the theoretical model for which the key factor defining the on-surface aggregation process is transient binding of monomers that play the role as nuclei in the assembly of aggregates. Importantly, the model works not only for α-Syn, a typical member of amyloidogenic proteins capable of assembly of into fibrils, as supported by numerous studies including ours (21-23), but also for A3G enzyme, for which the stoichiometry of aggregates is defined by the concentration of monomers (20). Notably, unlike the amyloid aggregates that dramatically change the physiological function of monomers ((24) and references therein), the assembly of A3G into oligomers contributes to its anti-HIV activity of A3G ((25) and references therein).

According top the theory (Eq. 2), the kinetics of on-surface aggregation depends on the affinity of the protein to the surface. The elevated propensity of A3G compared with α-Syn to form aggregates on surface points to its high affinity for the mica surface; this feature is in line with the high affinity of this protein to cellular the membrane and other intracellular particles (26, 27).

Although the experimental results support the theoretical prediction regarding the linear dependence for the initial aggregation process on time (Eq. 13), the dependence on the concentration is not fully in line with theoretical predictions. According to Table S1, the five-fold increase in the α-Syn concentration does not change the aggregation rate five-fold; only a two-fold increase is observed. However, this result is not surprising considering the theory does not include factors such as changes in the protein conformation upon the interaction with surfaces. Moreover, the formation of oligomers larger than dimers also can influence aggregation kinetics. Indeed, our computational simulation previously showed that amyloid beta peptide (Aβ) (14-23) undergoes a conformational change that facilitates the assembly of dimers (17). Based on our recent studies for the aggregation of α-Syn aggregation on membrane surfaces (28), it is reasonable to assume that α-Syn also undergoes a conformational change at the mica-liquid interface. Meanwhile, comparison of α-Syn aggregation on membrane bilayers of different compositions induced different conformational changes and resulted in only several-fold changes to the aggregation propensities of membranes (28). This value is considerably less than the overall aggregation catalysis of membranes and mica, which is in the range of several orders of magnitude (17); such an acceleration is in line with the current theoretical predictions. However, understanding the effect of the membrane composition on the entire on-surface aggregation process can explain the role of membrane surfaces in the assembly of amyloid aggregates (28) and will help elucidate the molecular mechanisms behind protein-aggregation diseases such as Alzheimer’s and Parkinson’s. The development of such a more-comprehensive model is our long-term goal.

## Supporting Information

Supporting methods, eight figures, and one table.

## Author Contributions

YLL and ABK designed the project. YP, SB, KZ, and LSS performed AFM experiments and data analysis. ABK provided the theoretical model. All authors contributed to writing the manuscript.

## Acknowledgments

This work was supported by grants to YLL from NIH (R01-GM118006, GM096039, and R21NS101504). ABK was supported by the Welch Foundation (C-1559), the NSF (CHE-1664218), and the Center of Theoretical Biological Physics, sponsored by the NSF (PHY-1427654). We thank Dr. R. Harris and Dr. C. Rochet for providing A3G and a-Syn proteins, respectively, and Dr. Lyubchenko lab members for their fruitful discussions. We thank Melody A. Montgomery for editing this manuscript.

## References

1. Hendrickson, W. A., A. Pahler, J. L. Smith, Y. Satow, E. A. Merritt, and R. P. Phizackerley. 1989. Crystal structure of core streptavidin determined from multiwavelength anomalous diffraction of synchrotron radiation. Proc Natl Acad Sci U S A 86(7):2190–2194.

2. Tamulaitis, G., M. Rutkauskas, M. Zaremba, S. Grazulis, G. Tamulaitiene, and V. Siksnys. 2015. Functional significance of protein assemblies predicted by the crystal structure of the restriction endonuclease BsaWI. Nucleic Acids Res 43(16):8100–8110.

3. Hardy, J., and B. De Strooper. 2017. Alzheimer’s disease: where next for anti-amyloid therapies? Brain 140(4):853–855.

4. Gray, J. J. 2004. The interaction of proteins with solid surfaces. Curr Opin Struct Biol 14(1):110–115.

5. Kurnik, M., G. Ortega, P. Dauphin-Ducharme, H. Li, A. Caceres, and K. W. Plaxco. 2018. Quantitative measurements of protein-surface interaction thermodynamics. Proc Natl Acad Sci U S A 115(33):8352–8357.

6. Andreasen, M., N. Lorenzen, and D. Otzen. 2015. Interactions between misfolded protein oligomers and membranes: A central topic in neurodegenerative diseases? Biochim Biophys Acta 1848(9):1897–1907.

7. Lindberg, D. J., E. Wesén, J. Björkeroth, S. Rocha, and E. K. Esbjörner. 2017. Lipid membranes catalyse the fibril formation of the amyloid-β (1–42) peptide through lipid-fibril interactions that reinforce secondary pathways. Biochimica et Biophysica Acta (BBA) - Biomembranes 1859(10):1921–1929.

8. Williams, T. L., and L. C. Serpell. 2011. Membrane and surface interactions of Alzheimer’s Abeta peptide--insights into the mechanism of cytotoxicity. FEBS J 278(20):3905–3917.

9. Canale, C., R. Oropesa-Nuñez, A. Diaspro, and S. Dante. 2017. Amyloid and membrane complexity: The toxic interplay revealed by AFM. Seminars in Cell & Developmental Biology 73: 82–94.

10. Bucciantini, M., S. Rigacci, and M. Stefani. 2014. Amyloid Aggregation: Role of Biological Membranes and the Aggregate-Membrane System. J Phys Chem Lett 5(3):517–527.

11. Roher, A. E., T. A. Kokjohn, S. G. Clarke, M. R. Sierks, C. L. Maarouf, G. E. Serrano, M. S. Sabbagh, and T. G. Beach. 2017. APP/Aβ structural diversity and Alzheimer’s disease pathogenesis. Neurochemistry International 110(Supplement C):1–13.

12. Cheng, B., Y. Li, L. Ma, Z. Wang, R. B. Petersen, L. Zheng, Y. Chen, and K. Huang. 2018. Interaction between amyloidogenic proteins and biomembranes in protein misfolding diseases: Mechanisms, contributors, and therapy. Biochim Biophys Acta 1860, 1876–88.

13. Rabe, M., A. Soragni, N. P. Reynolds, D. Verdes, E. Liverani, R. Riek, and S. Seeger. 2013. On-surface aggregation of alpha-synuclein at nanomolar concentrations results in two distinct growth mechanisms. ACS Chem Neurosci 4(3):408–417.

14. Lin, Y. C., C. Li, and Z. Fakhraai. 2018. Kinetics of Surface-Mediated Fibrillization of Amyloid-beta (12-28) Peptides. Langmuir 34(15):4665–4672.

15. Lin, Y. C., E. J. Petersson, and Z. Fakhraai. 2014. Surface effects mediate self-assembly of amyloid-beta peptides. ACS Nano 8(10):10178–10186.

16. Lucas, M. J., and B. K. Keitz. 2018. Influence of Zeolites on Amyloid-β Aggregation. Langmuir 34(33):9789–9797.

17. Banerjee, S., M. Hashemi, Z. Lv, S. Maity, J. C. Rochet, and Y. L. Lyubchenko. 2017. A novel pathway for amyloids self-assembly in aggregates at nanomolar concentration mediated by the interaction with surfaces. Sci Rep 7:45592–602.

18. Proctor, E. A., L. Fee, Y. Tao, R. L. Redler, J. M. Fay, Y. Zhang, Z. Lv, I. P. Mercer, M. Deshmukh, Y. L. Lyubchenko, and N. V. Dokholyan. 2016. Nonnative SOD1 trimer is toxic to motor neurons in a model of amyotrophic lateral sclerosis. Proc Natl Acad Sci U S A 113(3):614–619.

19. Banerjee, S., Z. Sun, E. Y. Hayden, D. B. Teplow, and Y. L. Lyubchenko. 2017. Nanoscale Dynamics of Amyloid beta-42 Oligomers As Revealed by High-Speed Atomic Force Microscopy. ACS Nano 11(12):12202–12209.

20. Shlyakhtenko, L. S., A. Y. Lushnikov, A. Miyagi, M. Li, R. S. Harris, and Y. L. Lyubchenko. 2013. Atomic force microscopy studies of APOBEC3G oligomerization and dynamics. J Struct Biol 184(2):217–225.

21. McAllister, C., M. A. Karymov, Y. Kawano, A. Y. Lushnikov, A. Mikheikin, V. N. Uversky, and Y. L. Lyubchenko. 2005. Protein Interactions and Misfolding Analyzed by AFM Force Spectroscopy. J Mol Biol 354(5):1028–1042.

22. Lyubchenko, Y. L., S. Sherman, L. S. Shlyakhtenko, and V. N. Uversky. 2006. Nanoimaging for protein misfolding and related diseases. J Cell Biochem 99:53–70.

23. Yu, J., S. Malkova, and Y. L. Lyubchenko. 2008. alpha-Synuclein misfolding: single molecule AFM force spectroscopy study. J Mol Biol 384(4):992–1001.

24. Copani, A. 2017. The underexplored question of β-amyloid monomers. European Journal of Pharmacology 817(Supplement C):71–75.

25. Shlyakhtenko, L. S., S. Dutta, J. Banga, M. Li, R. S. Harris, and Y. L. Lyubchenko. 2015. APOBEC3G Interacts with ssDNA by Two Modes: AFM Studies. Sci Rep 5:15648–59.

26. Moris, A., S. Murray, and S. Cardinaud. 2014. AID and APOBECs span the gap between innate and adaptive immunity. Front Microbiol 5:534–46.

27. Posch, W., M. Steger, U. Knackmuss, M. Blatzer, H. M. Baldauf, W. Doppler, T. E. White, P. Hortnagl, F. Diaz-Griffero, C. Lass-Florl, H. Hackl, A. Moris, O. T. Keppler, and D. Wilflingseder. 2015. Complement-Opsonized HIV-1 Overcomes Restriction in Dendritic Cells. PLoS Pathog 11(6):e1005005–27.

28. Lv, Z., M. Hashemi, S. Banerjee, K. Zagorski, J.-C. Rochet, and Y. L. Lyubchenko. 2018. Phospholipid membranes promote the early stage assembly of a-synuclein aggregates. bioRxiv preprint.

